# β-catenin-mediated activation of Wnt target genes utilizes a biomolecular condensate-dependent mechanism

**DOI:** 10.1101/2023.10.09.561634

**Authors:** Richard A. Stewart, Lauren B. Goodman, Jeannine J. Tran, John P. Zientko, Malavika Sabu, Ung Seop Jeon, Kenneth M. Cadigan

## Abstract

The Wnt/β-catenin signaling pathway plays numerous, essential roles in animal development and tissue/stem cell maintenance. The activation of genes regulated by Wnt/β-catenin signaling requires the nuclear accumulation of β-catenin, a transcriptional co-activator. β-catenin is recruited to many Wnt-regulated enhancers through direct binding to T-cell factor/Lymphoid enhancer factor (TCF/LEF) family transcription factors. β-catenin has previously been reported to form phase-separated biomolecular condensates (BMCs), which was implicated as a component of β-catenin’s mechanism of action. This function required aromatic amino acid residues in the intrinsically disordered regions (IDRs) at the N- and C-termini of the protein. In this report, we further explore a role for β-catenin BMCs in Wnt target gene regulation. We find that β-catenin BMCs are miscible with LEF1 BMCs *in vitro*. We characterized a panel of β-catenin mutants with different combinations of aromatic residue mutations in human cell culture and *Drosophila melanogaster*. Our data support a model in which aromatic residues across both IDRs contribute to BMC formation *in vitro* and signaling activity *in vivo*. Although different Wnt targets have different sensitivities to loss of β-catenin’s aromatic residues, the activation of every target examined was compromised by aromatic substitution. These mutants are not defective in nuclear import, and residues in the N-terminal IDR with no previously known role in signaling are clearly required for the activation of various Wnt readouts. Consistent with this, deletion of the N-terminal IDR results in a loss of signaling activity, which can be rescued by the addition of heterologous IDRs enriched in aromatic residues. Overall, our work supports a model in which the ability of β-catenin to form biomolecular condensates in the nucleus is tightly linked to its function as a transcriptional co-regulator.

## Introduction

The Wnt/β-catenin signaling pathway is evolutionarily conserved across metazoans and is indispensable for organismal development and a variety of adult tissue functions [1,2]. The primary output of this pathway is the differential regulation of gene expression programs, which is accomplished through the nuclear accumulation of β-catenin, a transcriptional co-regulator. β-catenin regulates Wnt targets in conjunction with transcription factors, the most prominent of which are members of the T-cell factor/lymphoid enhancer binding factor (TCF/LEF) family. Nuclear β-catenin binds with TCF/LEFs on the chromatin at *cis*-regulatory Wnt-responsive enhancers (WREs) [3,4]. Many cancers are causally linked to the inappropriate elevation of nuclear β-catenin, which can occur through loss of function mutations in negative regulators of the pathway, such as Adenomatosis polyposis coli (APC), Axin, and Ring finger protein 43 (RNF43), or through oncogenic mutations in β-catenin that prevent its turnover [5].

β-catenin is conventionally understood to be comprised of three distinct domains: an intrinsically disordered N-terminal domain (N-IDR), a highly structured internal domain consisting of twelve Armadillo (Arm) repeats, followed by an intrinsically disordered C-terminal domain (C-IDR) [3,6]. β-catenin’s N-IDR is necessary for regulating the stability of the protein and contains a region bound by α-catenin [6]. Cytosolic β-catenin is bound by a “destruction complex”, which contains APC, Axin, as well as two kinases, Casein Kinase I (CKI) and Glucagon Synthase Kinase-3 (GSK3), which serially phosphorylate four serine/threonine residues, priming the protein for proteasomal degradation [7]. The Arm repeat region contains binding sites for TCF/LEF transcription factors and E-cadherin (repeats 3-8), as well as co-activators BCL9 and BCL9L (repeat 1) [3,8–10]. The last four Arm repeats and the C-IDR are bound by a variety of transcriptional regulators, including chromatin remodelers such as Brg-1 and the histone acetyltransferases CBP/p300 [3,11–13]. The current model suggests that factors are sequentially recruited to WRE chromatin (i.e. TCF/LEFs recruit β-catenin, β-catenin recruits additional co-regulators) and this is sometimes referred to as the “chain of adaptors” model [14]. While there is significant support for TCF/LEFs, β-catenin, and other co-regulators physically interacting with each other to promote transcription, the exact nature of these interactions remains to be determined.

Recent studies indicate a role for biomolecular condensates (BMCs) in transcriptional activation [15–18]. BMCs are dynamic, membraneless assemblies comprised of proteins and, frequently, nucleic acids. Weak, multivalent interactions between the IDRs of constituent proteins drive the formation of BMCs, which is usually thought to occur through a liquid-liquid phase separation mechanism. Functionally, BMCs are thought to affect biochemical reactions by concentrating molecules, which can have potentiating or inhibitory effects [15]. The evidence for BMCs having a role in transcriptional regulation is derived from live imaging studies demonstrating the existence of dynamic puncta at enhancer chromatin and the propensity for many transcriptional regulators to form BMCs *in vitro* [15,17,19]. However, rigorous evidence for a physiological role for BMCs, e.g., provided by specific mutations in transcriptional regulators is still lacking.

A previous report by Zamudio and colleagues demonstrated that β-catenin protein can form homotypic and heterotypic BMCs *in vitro* [20]. β-catenin’s terminal IDRs were necessary and sufficient for BMC formation, and this behavior was dependent on aromatic amino acid residues within the IDRs. An endogenously GFP-tagged β-catenin was shown to form dynamic puncta in response to Wnt signaling in cultured cells, providing evidence that β-catenin-containing BMCs exist in living cells. Consistent with this, a mutant of β-catenin with IDRs lacking aromatic residues (19 total aromatic amino acid substitutions in the terminal IDRs) was defective in regulating Wnt target genes, recruitment to WRE chromatin, and puncta formation [20].

While the work of Zamudio and colleagues is consistent with a physiological role for β-catenin condensates, several key issues remain uncertain. For example, it was not clear in that report that condensate-deficient β-catenin accumulates in the nucleus at levels comparable to wild-type β-catenin. Additionally, the alteration of 19 aromatic residues in β-catenin’s terminal IDRs may disrupt key protein-protein interactions that are essential for β-catenin function, regardless of a condensate mechanism. These factors are potential explanations for the defect in recruitment to chromatin and transcriptional activation of the β-catenin aromatic mutant in that report [20]. Whether the ability of β-catenin to form BMCs is linked to its activity as a transcriptional co-activator requires further investigation.

In this report, we address the hypothesis that BMC formation is important for β-catenin-mediated regulation of Wnt target genes by generating and characterizing a panel of β-catenin mutants utilizing *in vitro* and *in vivo* experimental systems. The results support a model in which the aromatic amino acid residues in both the N- and C-IDRs contribute to BMC formation and transcriptional activity. Importantly, we found that the N-IDR of β-catenin has a previously underappreciated role in transcriptional regulation [21]. Supporting these findings, β-catenin was found to efficiently form heterotypic condensates with LEF1 *in vitro*, which also depends on IDR aromaticity. Transgenic *Drosophila* lines expressing analogous Armadillo (Arm, the fly ortholog of β-catenin) mutants demonstrated the importance of aromatic residues for Arm activity in the context of fly development. Interestingly, while the mutants displayed lower signaling activity, different Wnt targets (in both human cells and *Drosophila*) had different sensitivities to the loss aromatic residues. Finally, heterologous IDRs from proteins with no known role in transcriptional regulation were able to functionally replace the N-IDR of β-catenin, providing compelling evidence that the ability of β-catenin to form biomolecular condensates is inextricably linked to its role as a transcriptional activator of Wnt target genes.

## Results

### Aromatic amino acid residues within β-catenin’s terminal IDRs contribute to homotypic and heterotypic condensate formation *in vitro*

For many proteins, BMC formation is thought to arise from weak, multivalent interactions between protein IDRs [15]. The N- and C-terminal regions of β-catenin are predicted to be disordered, illustrated by two independent methods of analysis, IUPred2A [22,23] and AlphaFold [24,25] (S1A and S1B Fig). Previous work from Zamudio and colleagues demonstrated that β-catenin can form homotypic BMCs *in vitro*, and that the terminal IDRs are necessary and sufficient for condensate formation. Additionally, they showed that the aromatic amino acid residues within both IDRs are required for this behavior [20]. However, the contributions of aromatic residues from the individual IDRs were not examined. Given that these regions have distinct roles in β-catenin function, i.e., N-IDR contains phosphorylation sites controlling β-catenin stability and C-IDR is required for co-regulator activity, we were motivated to examine the requirements for the aromatic residues in more detail.

To address the role of aromatics in each IDR of β-catenin, we recombinantly expressed four eGFP-β-catenin fusion proteins: one containing the full set of aromatic residues plus a S33Y point mutation (β-catenin*). This mutation was incorporated for direct comparison with the β-catenin mutants used for subsequent functional studies. Additional proteins have the N-IDR aromatic residues mutated to alanine (aroN; 9 substitutions), the C-IDR aromatics mutated to alanine (aroC; 10 substitutions), and aromatics in both IDRs mutated to alanine (aroNC; 19 substitutions) (Fig 1A and S2 Fig for sequence information). These proteins were tested for condensate formation using a buffer containing 10% polyethylene glycol 8000 (PEG-8000) as a crowding agent, following standard experimental guidelines [26]. At relatively low concentrations where β-catenin* efficiently formed droplets, the aromatic mutants are deficient in condensate formation (Fig 1B). aroN and aroC displayed a similar defect in droplet formation, while aroNC was more severe. At high concentrations, the aromatic mutants formed condensates at similar levels as β-catenin*. Our results with aroNC are inconsistent with those reported by Zamudio and colleagues (see Discussion for further comment). However, consistent with that report, eGFP alone or eGFP-β-catenin with both IDRs deleted (ΔNC; Fig 1A) is incapable of condensate formation (Fig 1B). Our results demonstrate that both the N-IDR and C-IDR contribute to β-catenin condensation, and these IDRs contain additional sequence information beyond aromatic residues that facilitate droplet formation.

**Fig 1.**
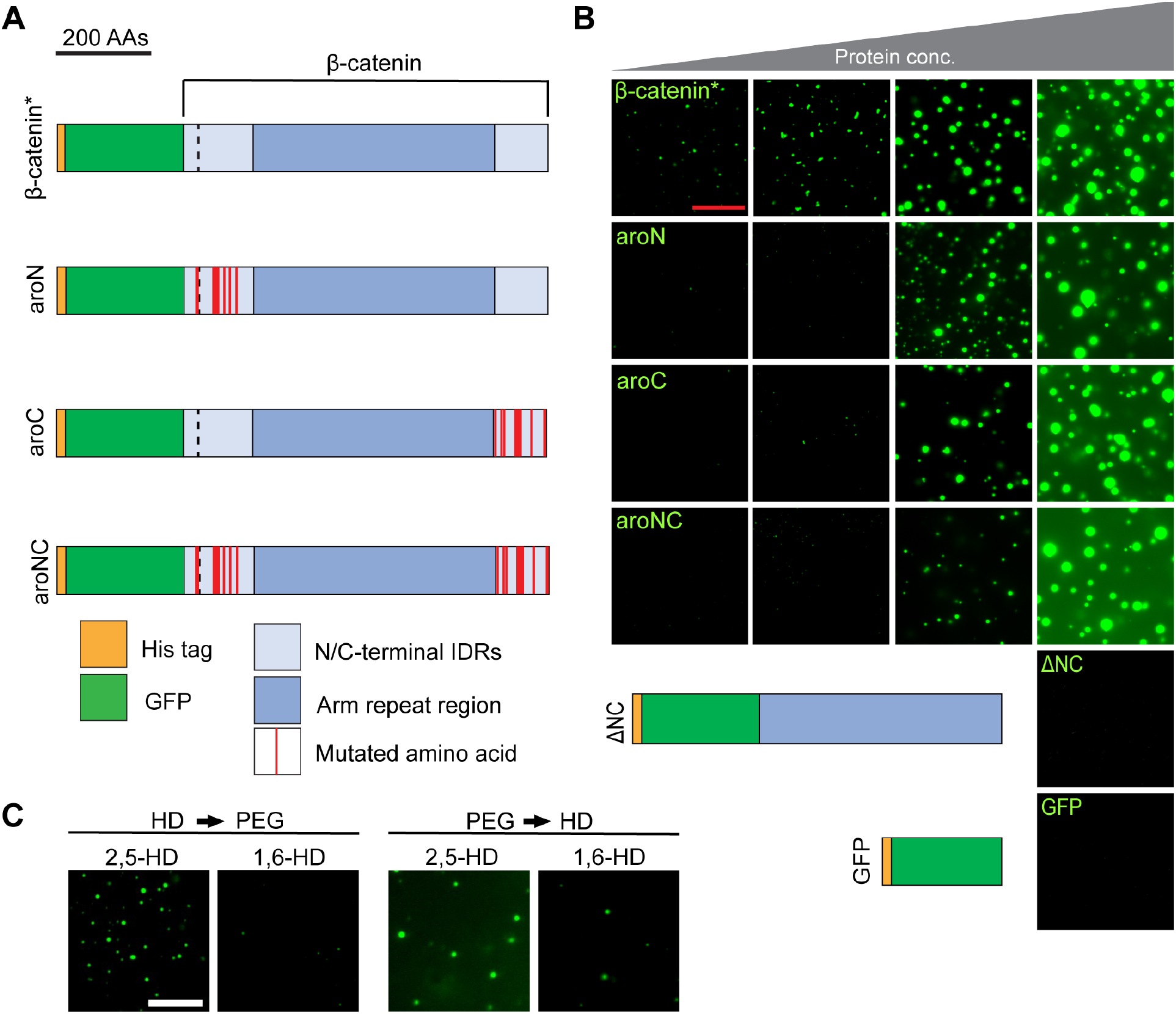
Aromatic amino acid residues in the terminal IDRs of β-catenin promote biomolecular condensate formation *in vitro*. (A) Cartoon representation of the eGFP-β-catenin* protein, the aromatic mutant derivatives, and control constructs. β-catenin* contains one S33Y mutation (dashed black line) in the N-IDR. aroN has all 9 endogenous aromatic amino acids within the N-IDR mutated to alanines. aroC has all 10 endogenous aromatic amino acids within the C-IDR mutated to alanines. aroNC contains all 19 aromatic amino acid mutations. The aromatic mutant constructs contain the same S33Y mutation as β-catenin*. Representative images from an *in vitro* droplet formation assay with the indicated mutants. A protein concentration series of 2μM, 3μM, 4μM, and 8μM is depicted. Droplet assays were performed in 300mM NaCl and 10% PEG-8000. Scale bar = 20μm. (C) Representative images from an *in vitro* droplet formation assay testing eGFP-β-catenin* sensitivity to 1,6-hexanediol and 2,5-hexanediol (used as a control). 8μM eGFP-β-catenin* protein was exposed to the hexanediols prior to PEG-8000 (HD -> PEG) or PEG-8000 prior to the hexanediols (PEG -> HD). 10% hexanediol and 10% PEG-8000 was used. Scale bar = 20μm.

To further examine the properties of the β-catenin* condensates we generated, we made use of the alcohol 1,6-hexanediol (1,6-HD), which is commonly used to inhibit biomolecular condensation [27,28]. 2,5-hexanediol (2,5-HD), which is chemically similar but doesn’t disrupt condensation was used as a control. Droplet formation of β-catenin* was sensitive to 1,6-HD, especially when the alcohol was added prior to the PEG-8000 crowding agent (Fig 1C). When 1,6-HD was added after PEG-8000, some condensation was still observed. The sensitivity to 1,6-HD is consistent with the idea that these assemblies are driven by hydrophobic interactions [28–30], but once the droplets form, some may transition from a phase separated droplet to more static hydrogel, which can resist the effects of 1,6-HD [31].

In living cells, β-catenin regulates gene expression through interactions with many proteins, the most prominent of which are members of the TCF/LEF family of transcription factors [3]. The central Arm repeats of β-catenin bind to the N-terminus of TCFs, as shown by traditional protein interaction assays and X-ray diffraction of co-crystals [10]. To determine if β-catenin could form heterotypic *in vitro* condensates with a TCF/LEF member, we expressed human mCherry fused to LEF1 (Fig 2A and S3A Fig for sequence information). IUPred2A and AlphaFold analysis of LEF1 predicts that most of the protein is disordered, except for the DNA-binding HMG domain (S1C and S1D Fig). Purified mCherry-LEF1 can form concentration-dependent droplets (Fig 2B). Next, we performed a dose with equal molar amounts of β-catenin* and LEF1 (Fig 2C). There is a high degree of co-localized fluorescent signal from both eGFP and mCherry, indicating a high degree of miscibility of β-catenin* and LEF1 droplets. Relative to β-catenin*, aroNC exhibits reduced co-localization with LEF1 (Fig 2D), as does ΔNC (Fig 2E). The degree of co-localization in individual BMCs was quantified with line traces, and as expected, the strongest colocalization was with β-catenin* and LEF1, followed by aroNC and ΔNC (Fig 2F-H and S4 Fig). Control heterotypic *in vitro* droplet formation assays show that eGFP does not co-localize with mCherry-LEF1 (S5A Fig) and mCherry does not co-localize with eGFP-β-catenin* (S5B Fig). The results indicate that β-catenin is localized to LEF1 condensates by two primary interactions. The traditional Arm repeat-LEF1 interaction makes a detectable contribution, based on the ability of LEF1 to recruit ΔNC into a mixed condensate (Fig 2E). However, the presence of the terminal IDRs greatly enhanced the ability of β-catenin and LEF1 to form mixed condensates (Fig 2C) and the aromatic residues in these IDRs are required (Fig 2D).

**Fig 2.**
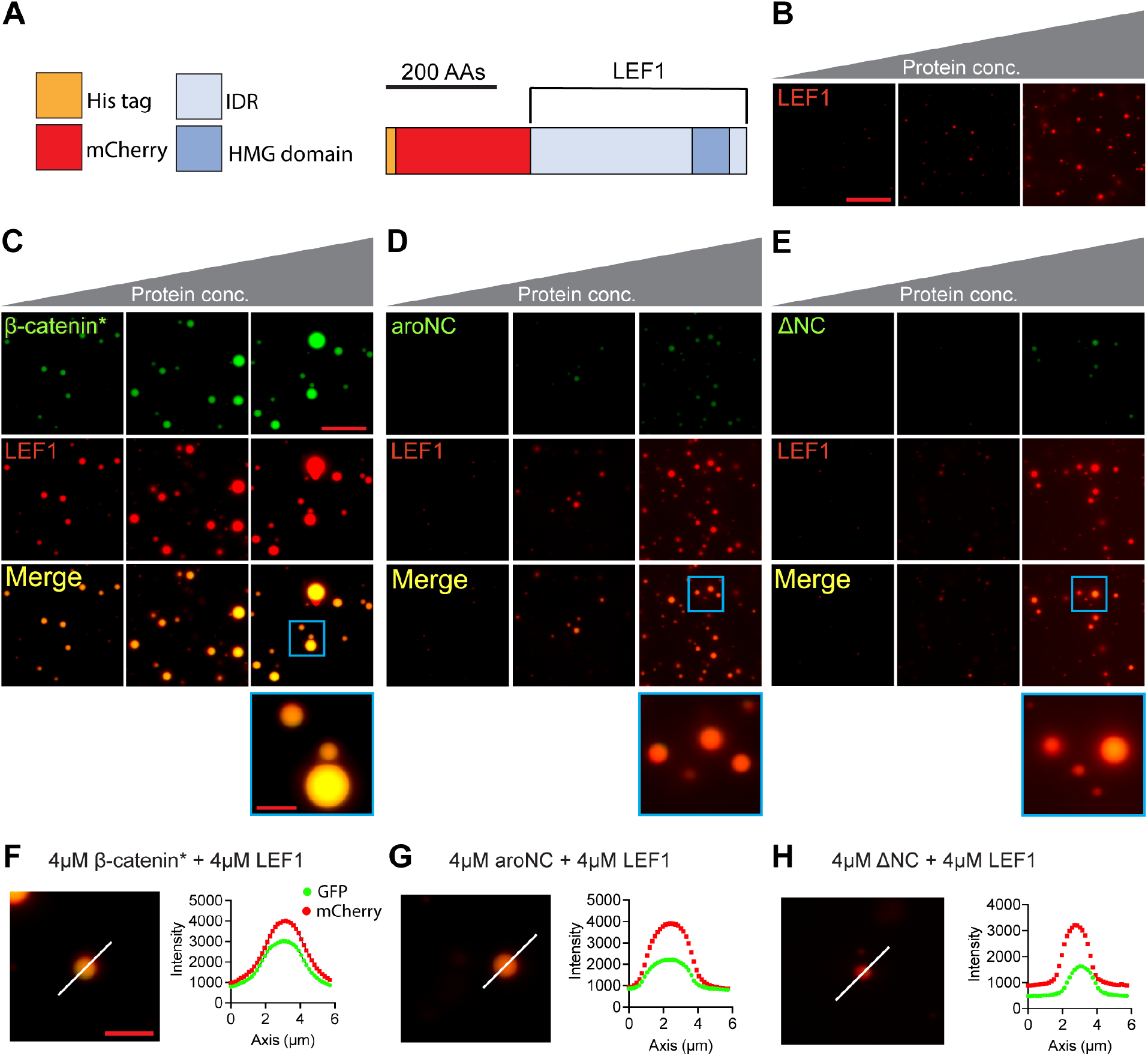
The β-catenin terminal IDRs promote β-catenin incorporation into Lef1 condensates *in vitro*. (A) Cartoon representation of the mCherry-LEF1 construct. (B) Representative images from an mCherry-LEF1 *in vitro* droplet formation assay. A concentration series of 4μM, 8μM, and 16μM total protein is depicted. Equal amounts of mCherry-LEF1 and eGFP (used as a control) were present in each reaction. No eGFP fluorescence was detected in the reaction and eGFP did not detectably affect the degree of mCherry-LEF1 droplet formation. Assays were performed in 300mM NaCl and 10% PEG-8000.Scale bar = 20μm. (C-E) Representative images from heterotypic *in vitro* droplet formation assays. (C) Equal amounts of eGFP-β-catenin* and mCherry-LEF1, (D) eGFP-aroNC and mCherry-LEF1, and (E) eGFP-ΔNC and mCherry-LEF1 were mixed, resulting in total protein concentrations of 4μM, 8μM, and 16μM. Reaction conditions are the same as panel B. Scale bar = 20μm, inset scale bar = 5μm. (F-H) Line plots showing eGFP-β-catenin* + mCherry-LEF1 (F), eGFP-aroNC and mCherry-LEF1 (G), and eGFP-ΔNC and mCherry-LEF1 (H) fluorescent intensity across a droplet. Increased fluorescent signal for both proteins across a line indicates co-localization and the white lines represent the plotted trace. Scale bar = 5μm.

### Aromatic amino acid residues within β-catenin’s IDRs are critical for nuclear function in cultured human cells

To test whether the ability of β-catenin to form homotypic and heterotypic condensates *in vitro* is relevant to its ability to activate Wnt targets in cultured cells, we expressed β-catenin*, aroN, aroC, and aroNC in HEK293T cells in the presence of several Wnt reporters. Using either the synthetic reporter TopFlash (containing six copies of high affinity TCF binding sites) [32] or a reporter with an endogenous WRE from the *Axin2* locus, known as CREAX [33], we found that all three β-catenin aromatic mutants had greatly reduced transcriptional activation activity (Fig 3A) even though they were expressed to similar levels as β-catenin* (Fig 3B). Similar results were also obtained with the Defa5-luc reporter that is synergistically activated by Wnt/β-catenin signaling and SOX9 (S6 Fig) [34]. In all three cases, aroN had some residual activity, aroC exhibited small but reproducible activity and aroNC had no detectable activity. To address whether the aromatic mutants were able to translocate into the nucleus, immunofluorescence (IF) was performed. We observed no detectable difference in the ability to these proteins to accumulate in the nucleus to similar levels as β-catenin* (Fig 3C and D). These results demonstrate the importance of the IDR aromatic residues for the ability of nuclear β-catenin to activate Wnt target gene expression. In addition, the defect in aroN reveals a previously unappreciated role for the N-terminus of β-catenin in transcriptional activation, which we suggest is due to the defect in the ability of the aromatic mutants to efficiently form BMCs.

**Figure 3.**
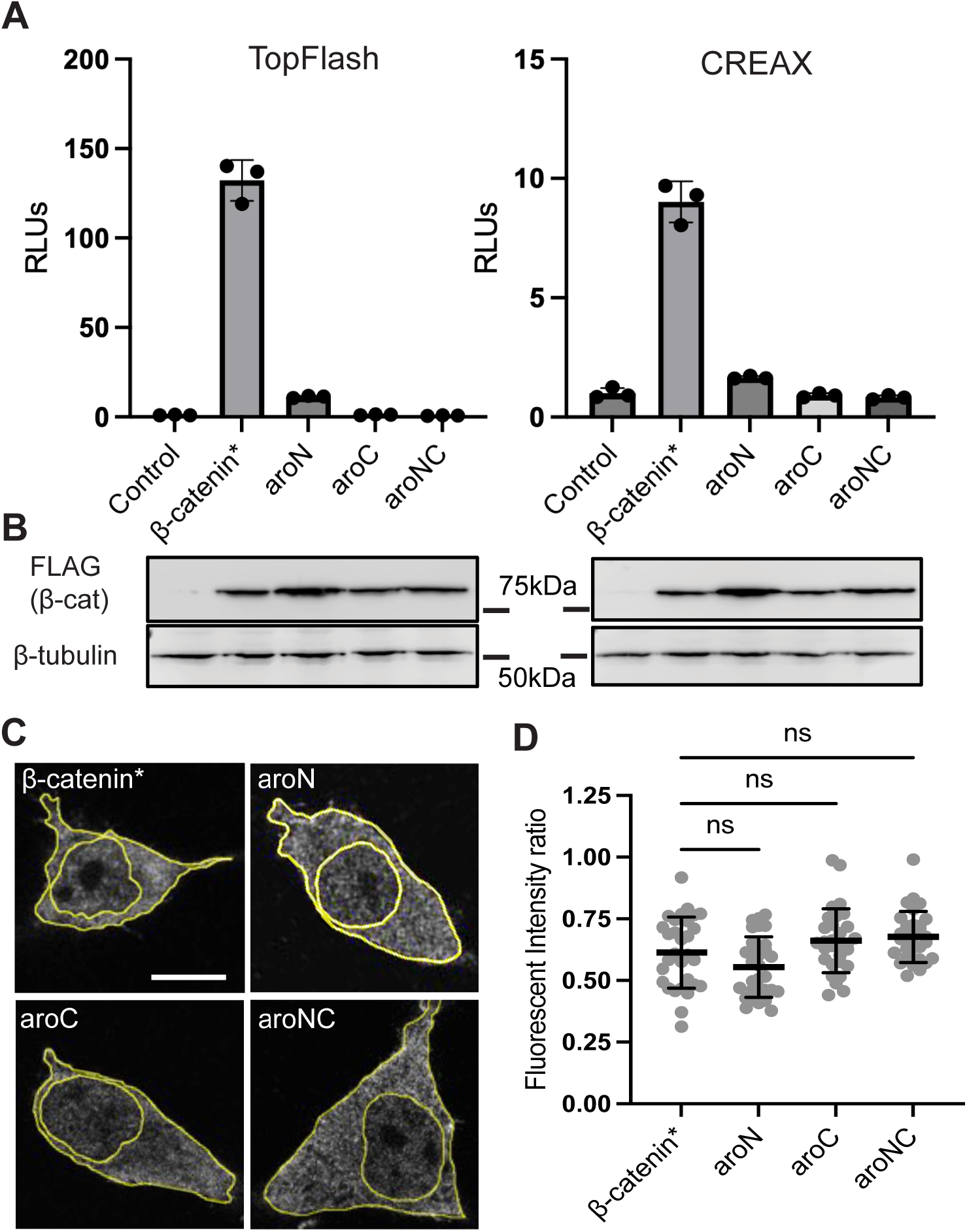
Aromatic residues within β-catenin’s terminal IDRs are required for reporter gene activation and not nuclear accumulation. (A) TopFlash (left) or CREAX (right) luciferase reporter activity induced by β-catenin* or the aromatic mutant constructs in HEK293T cells. Cells were transfected with separate plasmids encoding the reporter genes and the FLAG-β-catenin mutant constructs or pcDNA3.1 as a negative control (NC). Luciferase activity was assayed 24hr post-transfection. Data are plotted as mean ± SD (n = 3). (B) Western blots of the HEK293T lysates that were transfected with the luciferase reporter gene and the FLAG-β-catenin mutant constructs. The lysate samples correspond to the luciferase reporter assay. α-FLAG blot shows β-catenin expression, α-tubulin was used as a loading control. (C) Representative IF images of HEK293T cells for the indicated FLAG-β-catenin mutants. Cells were transfected with plasmids encoding the FLAG-β-catenin mutant constructs. IF was performed 24hr after transfection and the cells were stained with DAPI. The borders of the cell and nucleus are highlighted. Scale bar = 10μm. (D) Quantification of IF showing no significant difference in nuclear localization between β-catenin* and aromatic mutants. Data is presented as a ratio of the fluorescent intensity within the nucleus to the fluorescent intensity outside the nucleus. Data are presented as mean ± SD (n = 30). p-values were calculated by one-way ANOVA followed by Dunnett’s test. ns = p > 0.05.

Our studies clearly indicate that aromatic residues in both IDRs are necessary for signaling activity, but further mutagenesis is needed to determine whether all aromatic residues equally contribute to β-catenin activity. One popular model of condensation driven by aromatic residues posits that aromatic residues flanked by polar amino acids (i.e., stickers) are the drivers of IDR-IDR interactions, while other aromatics serve as “spacers” [35]. To test this sticker/spacer model in the context of β-catenin transcriptional activity, we constructed four additional mutants (S6D Fig for sequence information). Per the model, we mutated five potential stickers and four potential spacers in N-IDR, and five potential stickers and spacers each in the C-IDR. These mutants were tested for activity in the TopFlash reporter assay. All four mutants displayed reduced activity but were more active than their aroN or aroC counterparts (S6 Fig). While N-sticker had a two-fold greater reduction in activity than N-spacer, the putative spacer aromatics in the C-IDR were more critical for activity than the putative stickers. Overall, our results do not support a strict sticker/spacer model for β-catenin’s IDRs; the results are more consistent with a model where many/most of the aromatic residues in the IDRs contribute to biological activity.

To extend our analysis beyond reporter genes, we examined the role of the IDR aromatic residues in β-catenin’s ability to regulate endogenous Wnt targets. We generated stable HeLa cell lines which expressed β-catenin* and the aromatic mutants from a DOX-inducible expression cassette via lentiviral transduction. We chose to assay the Wnt target genes *Axin2* and *Sp5* as they are strongly activated by Wnt signaling in HeLa cells [36,37]. qPCR analysis of HeLa cells expressing β-catenin* and the aromatic mutants indicates that the relationship between aromatic amino acid residues and gene regulation is slightly more complex than the reporter activity (Fig 4A). For *Sp5*, all aromatic mutants are deficient, but not defective in activating expression. For *Axin2*, aroNC is defective in activating expression, while aroC is deficient and aroN has no detectable defect. These mutants were expression matched for comparison (Fig 4B). The observation that the aromatic mutants have a different effect on genes within the same cell type indicates that there can be gene-specific requirements for BMCs in Wnt target gene regulation. IF analysis revealed that β-catenin* forms puncta in these cells, while aroN is slightly deficient in forming puncta. aroC and aroNC do not form puncta (Fig 4C). These observations fit with the *in vitro* droplet formation data and roughly correlate with β-catenin’s ability to transcriptionally activate Wnt target genes.

**Fig 4.**
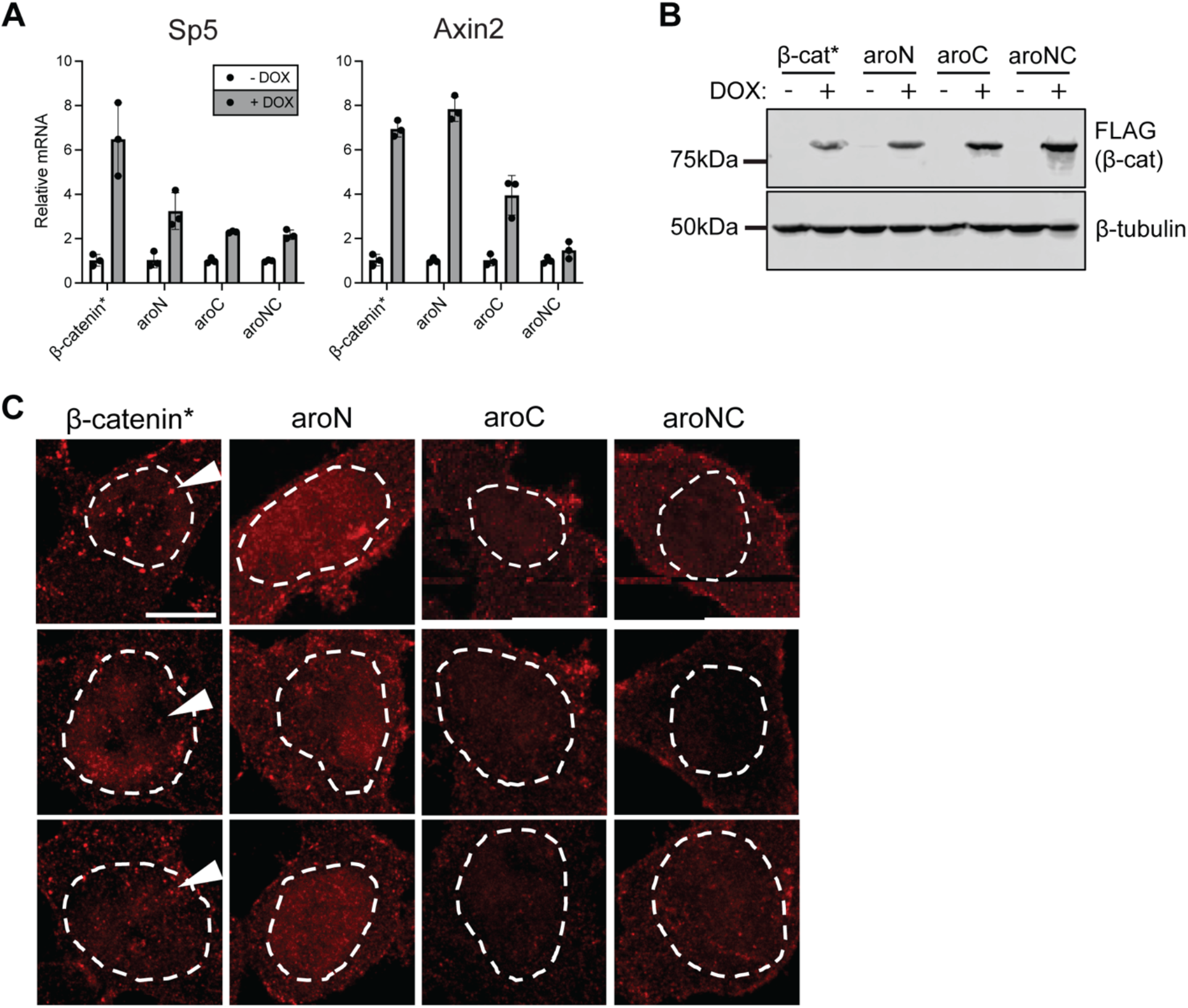
Select Wnt target genes exhibit different sensitivities to β-catenin aromatic mutant constructs. (A) qRT-PCR analysis of two Wnt target genes (*Sp5* and *Axin2*) in HeLa cells stably transformed with DOX-inducible, β-catenin mutant expression vectors. Cells were treated with DOX for 24hr. Data presented as mean ± SD (n=3). (B) Western blot analysis of DOX-treated HeLa cell lysate. Lysate samples correspond to the qPCR data. α-FLAG blot shows β-catenin expression. α-tubulin was used as a loading control. (C) Representative confocal IF images of HeLa cells expressing β-catenin mutants. 3 images per mutant are shown. Cells were treated with DOX for 24hr prior to IF. Cells were stained with DAPI, and dashed lines represent the nuclear border. Arrow indicates the presence of β-catenin* puncta, which are reduced in aroN expressing cells and not detectable in aroC and aroNC cells. Scale bar = 10μm.

### β-catenin/Armadillo IDR aromatic residues are critical for function in *Drosophila* development

To test whether the importance of aromatic residues for β-catenin function is conserved across species, we examined their role in Arm. Wingless (Wg)/Arm signaling is required throughout *Drosophila* development and has been intensively studied in *Drosophila* embryos and larval imaginal discs [38,39]. Similar to human β-catenin, Arm contains 9 and 10 aromatic residues in its N-IDR and C-IDR, respectively. Seven of the N-terminal aromatics and 5 of the C-terminal aromatics are conserved across multiple species. We constructed 5 Arm transgenes, under the control of the Gal4-UAS expression system (S7 Fig for protein sequences). These transgenes were integrated into two locations in the fly genome using phiC31 landing sites [40], ensuring similar levels of transcription. These UAS-transgenes were expressed in various tissues and their effect on Wg/Arm readouts were assayed.

It has been previously shown that expressing Wg agonists in the larval eye via the GMR-Gal4 driver results in smaller eyes due to increased apoptosis [41,42]. Using GMR-Gal4 to overexpress UAS-Arm*, a constitutively active mutant, we observed a reduction in adult eye size, and the loss of pigmentation and cone cells (Fig 5A-C). All the aromatic Arm mutants (which contain the stabilizing mutants of Arm*) exhibit minimal signaling activity in this assay, though they are expressed at similar levels, as detected by IF (Fig 5D). These data indicate that the aromatic residues within both IDRs are essential for Arm signaling activity in the developing *Drosophila* eye.

**Fig 5.**
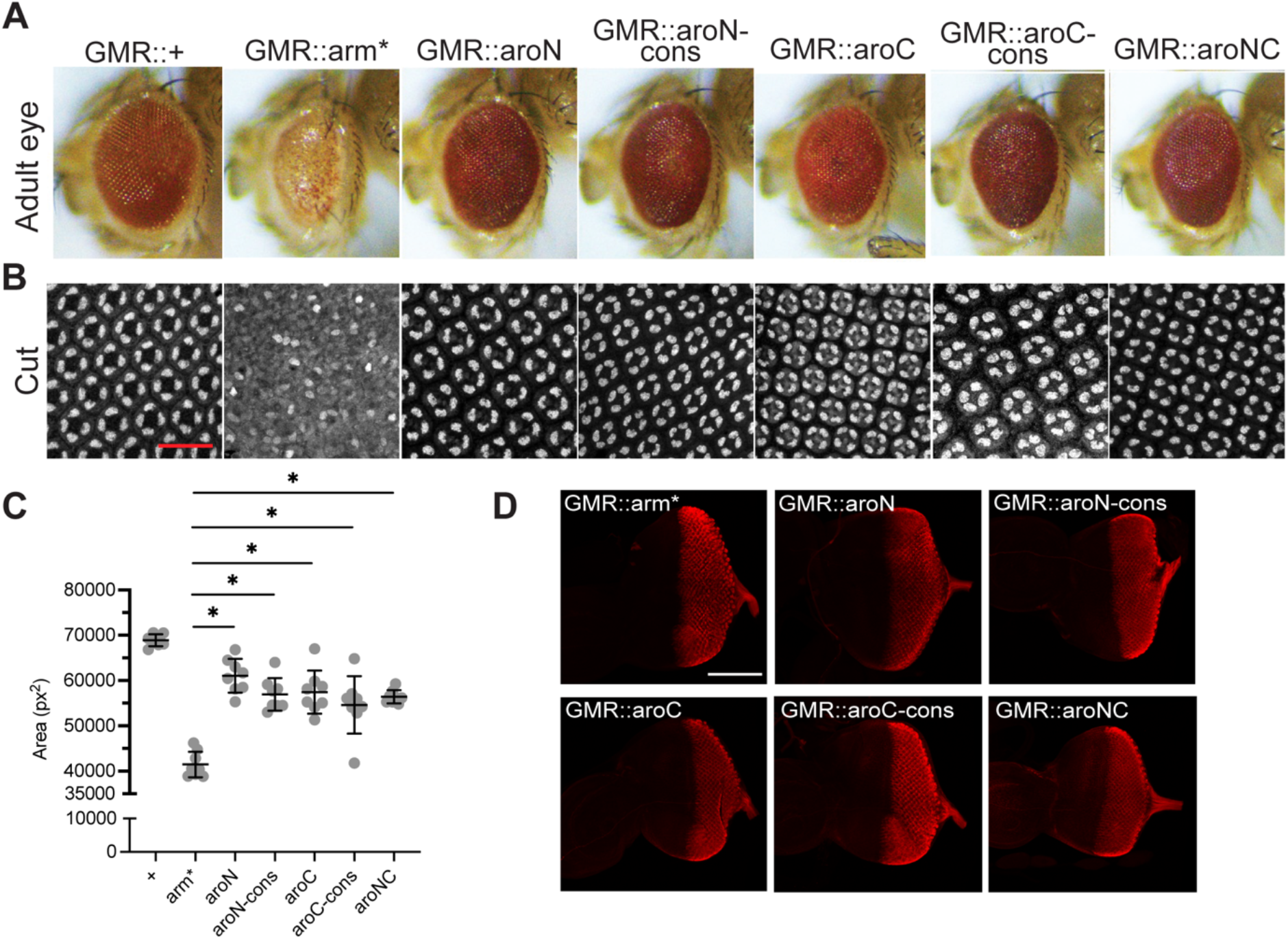
β-catenin/Arm activity in the adult Drosophila eye is attenuated by aromatic amino acid mutations within the terminal IDRs. (A) Micrographs of adult *Drosophila* eyes containing P[GMR-Gal4] and various P[UAS-Arm] transgenes. (B) Representative images of pupal *Drosophila* eye tissue immunostained for the cone cell marker Cut. Stabilized Arm (Arm*) disrupts cone cell specification while aromatic mutants do not. Scale bar = 20μm. (C) Quantification of adult *Drosophila* eye area. Data are presented as mean ± SD (n = 8). p-values were calculated by one-way ANOVA followed by Dunnett’s test. * = p < 0.05. (D) Expression of the various Arm constructs is constant during late larval eye development. Representative images of larval *Drosophila* eye antennal discs immunostained for FLAG, representing β-catenin mutant expression. Scale bar = 100μm.

Wg/Arm signaling is important for patterning the wing imaginal disc during larval development. In this tissue, Wg is expressed across the dorsal/ventral boundary in a stripe, regulating targets at short and long range from the source of Wg synthesis [38]. This gradient of signaling can be detected with a synthetic Wg reporter containing 4 Grainy head binding sites upstream of 4 HMG-Helper site pairs, arranged for high affinity binding by Pangolin (the fly TCF ortholog) [43–45]. Decapentaplegic-Gal4 (DPP-Gal4) was used to overexpress Arm* and the aromatic mutants in a stripe pattern that is perpendicular to the endogenous Wg expression stripe (Fig 6A, white arrowhead). To prevent major disruption to the wing disc morphology, the Gal80^ts^ system was used to inhibit Gal4 activity (and Arm protein expression) until 18 hours prior to fixation. Arm* overexpression resulted in the strongest ectopic activation of the Wg reporter, the aroN and aroN-cons constructs exhibited a moderate activation of the reporter, and the aroC, aroC-cons, and aroNC mutants exhibited the weakest ectopic activation (Fig 6A, top). The observations and categorizing the constructs Into strong/moderate/weak activators is supported by quantification of the fluorescent reporter activity and subsequent statistical analyses (S8 Fig). All mutant constructs were expressed at similar levels, as detected by FLAG IF (Fig 6A, bottom). These reporter data show that all the aromatic mutants have some residual capacity to activate transcription. These data are distinct from the developing eye, in which the aromatic mutants had little/no activity. This difference in sensitivity to loss of aromatic residues could be due to difference in the degree for BMC-dependency for activation of Wnt targets in different tissues.

**Fig 6.**
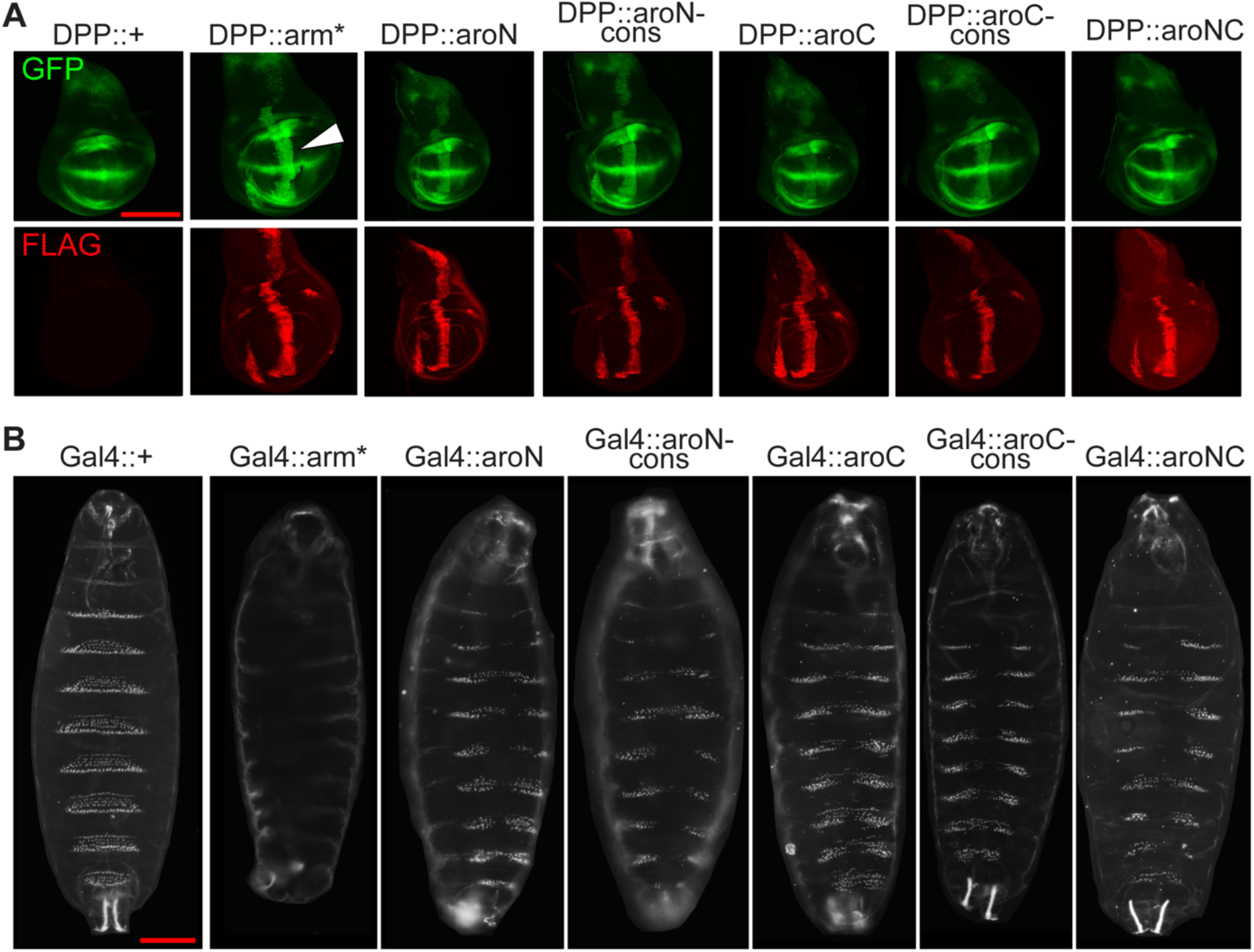
Aromatic β-catenin/Arm mutants exhibit different levels of activity in wing imaginal discs and embryonic epidermis. (A) Representative images of late third instar wing imaginal discs showing expression of a synthetic Wnt GFP reporter combined with a P[Dpp-Gal4] driving expression of various P[UAS-Arm] transgenes. Discs were also immunostained with α-FLAG to detect expression of the various Arm mutants. Gal4 activity was restricted to 18hr before fixation using a Gal80^ts^ transgene. Scale bar = 100μm. (B) Representative darkfield images showing ventral side of late embryonic *Drosophila* cuticles containing P[Da-Gal4], P[Arm-Gal4] and two copies of the various P[UAS-Arm] transgenes. Scale bar = 100μm.

Wg/Arm signaling also plays a key role in patterning the *Drosophila* embryo. Segments of the ventral embryonic epidermis feature a characteristic, trapezoidal-shaped belt of denticles. These denticle belts are separated by regions of naked cuticle. The establishment of denticle belts and naked cuticle is regulated by Wnt signaling [46]. Increasing Wnt signaling throughout the embryo expands regions of naked cuticle at the expense of denticle band formation, and conversely, loss of Wnt signaling leads to ectopic denticle formation and a failure to form naked cuticle [47]. To test the effect that our aromatic mutants have on regulating this phenotype, we overexpressed our constructs to similar levels using a stock containing two constitutive Gal4 drivers, Daughterless-Gal4 (Da-Gal4) and Arm-Gal4, both of which are active throughout the embryonic epidermis [48,49] (S9 Fig). When crossed to this Gal4 driver stock, UAS-Arm* displays a classic naked cuticle phenotype, with a 100% phenotype penetrance (Fig 6B). In contrast, overexpression of the aromatic mutants all resulted in a similar phenotype: a partial loss of denticle formation along the ventral midline (Fig 6B). These phenotypes were highly penetrant and consistent with a moderate level of Arm signaling activity. Our data indicates that the loss of aromatic residues in N-IDR and as little as five aromatic residues within the C-IDR (i.e., aroCcon) compromise Arm’s signaling activity to similar extents.

To test the ability of an Arm protein lacking IDR aromatic residues to rescue an *arm* loss-of-function phenotype, we expressed our transgenes at a reduced level (Da-Gal4 plus one copy of a UAS-Arm transgene). We reasoned that at this lower level of expression, the transgenic Arm* would be able to rescue the severe cuticular phenotype of embryos lacking zygotic *arm* gene activity. Indeed, expression of Arm* was able to rescue the *arm* mutant phenotype to a high-degree with 100% penetrance (Fig 7A-C). Expression of aroNC also resulted in significant rescue (Fig. 7D): the overall size of these embryos is similar to wild-type embryos and the Arm* rescue, head structures are restored and there is significant recovery of posterior-most structures. However, the degree of rescue was significantly less for aroNC than Arm*, as evidenced by the presence of excess denticles in all the abdominal segments. The difference in rescue may be reflected in the ability of these overexpression backgrounds to drive a naked cuticle phenotype in a wild-type Arm genetic background. Single-copy overexpression of Arm* causes a moderate naked cuticle phenotype, as some denticles are still present (Fig 7E), and aroNC overexpression exhibits weaker activity, as it only disrupts denticle formation in some segments (Fig 7F). The extent to which aroNC can rescue an *arm* loss of function phenotype relative to Arm* suggests a surprising degree of residual activity. Whether this is related to the residual ability of aroNC to form BMCs or because many molecular targets in embryonic epidermis do not require β-catenin condensation will require additional experiments (see Discussion for further comment).

**Fig 7.**
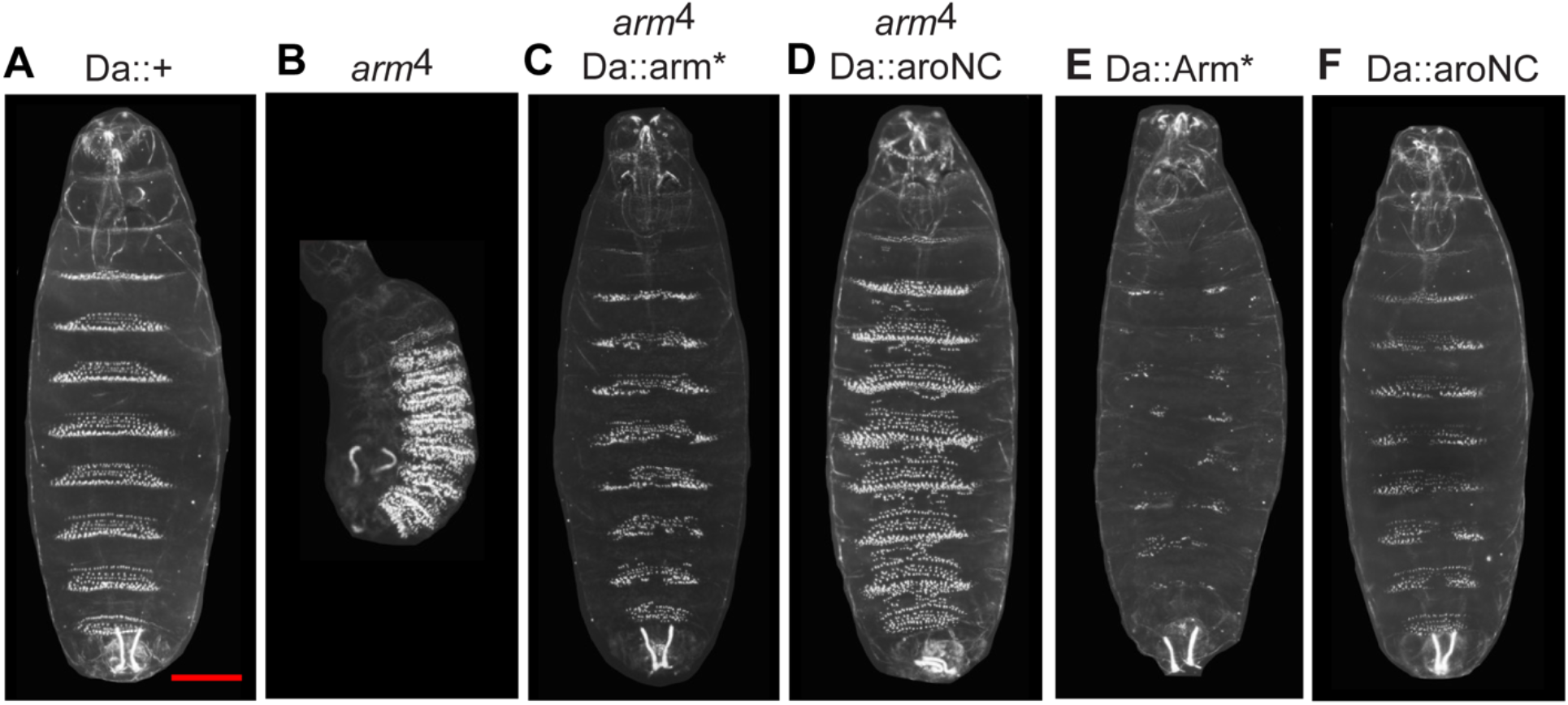
An aromatic β-catenin/Arm mutant (aroNC) partially rescues an *arm* loss-of-function allele. (A) Ventral side of a cuticle of an embryonic containing the P[Da-Gal4] transgene. Phenotype is indistinguishable from wild-type. (B) Cuticle of an amorphic *arm* mutant, displaying a classic Wg loss-of-function phenotype. The size of the embryo is greatly reduced, the head and posterior structures are missing or malformed, and the “naked” cuticle normally found on the posterior portion of each segment is absent, instead displaying ectopic denticles. (C) Cuticle of the *arm* mutant containing P[Da-Gal4] and P[UAS-Arm*]. This combination results in nearly complete rescue of the arm phenotype with 100% penetrance. (D) Cuticle of the *arm* mutant containing P[Da-Gal4] and P[UAS-aroNC]. The aromatic mutant rescues the size defect, most of the head structures, and some of the posterior. Segments still contain ectopic denticles indicating some reduction in Wg signaling. (E) Cuticle of an embryo containing P[Da-Gal4] and P[UAS-Arm*]. (F) Cuticle of an embryo containing P[Da-Gal4] and P[UAS-aroNC].

### Heterologous IDRs can rescue β-catenin signaling activity of a N-IDR deletion mutant

To this point, our mutagenesis approach has correlated a loss of aromatic residues within β-catenin’s IDRs with a loss of BMC formation and transcriptional regulation function, providing a link between the ability to form BMCs *in vitro* and function *in vivo*. This argument is problematic for the C-IDR, which has been implicated in binding to co-activators [50]. This caveat is mitigated by the fact that there are no known co-activator binding partners for N-IDR. As shown below, deletion of the N-IDR dramatically affected the ability of β-catenin to form droplets *in vitro* and activate some Wnt targets. This provided an opportunity to test whether these activities could be rescued by adding heterologous IDRs to β-catenin lacking the N-IDR. A collection of IDRs [51] was screened with the following criteria: (1) the IDR must come from a protein with no known nuclear function, (2) must be of similar size as N-IDR (∼140aa), and (3) must have a similar frequency of aromatic amino acids. Two IDRs, from human Septin 4 (Sept4) and Sorting nexin 18 (SNX18) met these criteria and were utilized (S10 Fig for protein sequences).

We generated eGFP-tagged β-catenin mutant constructs with the heterologous IDRs at the N-terminus (Fig 8A). We then performed a concentration series with the *in vitro* droplet formation assay (Fig 8B). Consistent with our previous observations, eGFP-β-catenin* will form spherical BMCs across the concentration range. In contrast, eGFP-ΔN forms fibril-like structures at relatively high concentrations. These fibril-like structures are morphologically distinct from the BMCs formed by the other mutants. eGFP-Sept4 and eGFP-SNX18 form BMCs that are similar in shape to β-catenin, rescuing the fibril-like structures of eGFP-ΔN. Furthermore, mutating the aromatic residues within the Sept4 IDR compromises the ability to form BMCs, reminiscent of the aroN mutant. These data suggest that an N-terminal IDR with sufficient aromatic content is required for BMC formation *in vitro*, and that the primary sequence of β-catenin’s endogenous N-IDR is not the driver of BMC formation.

**Fig 8.**
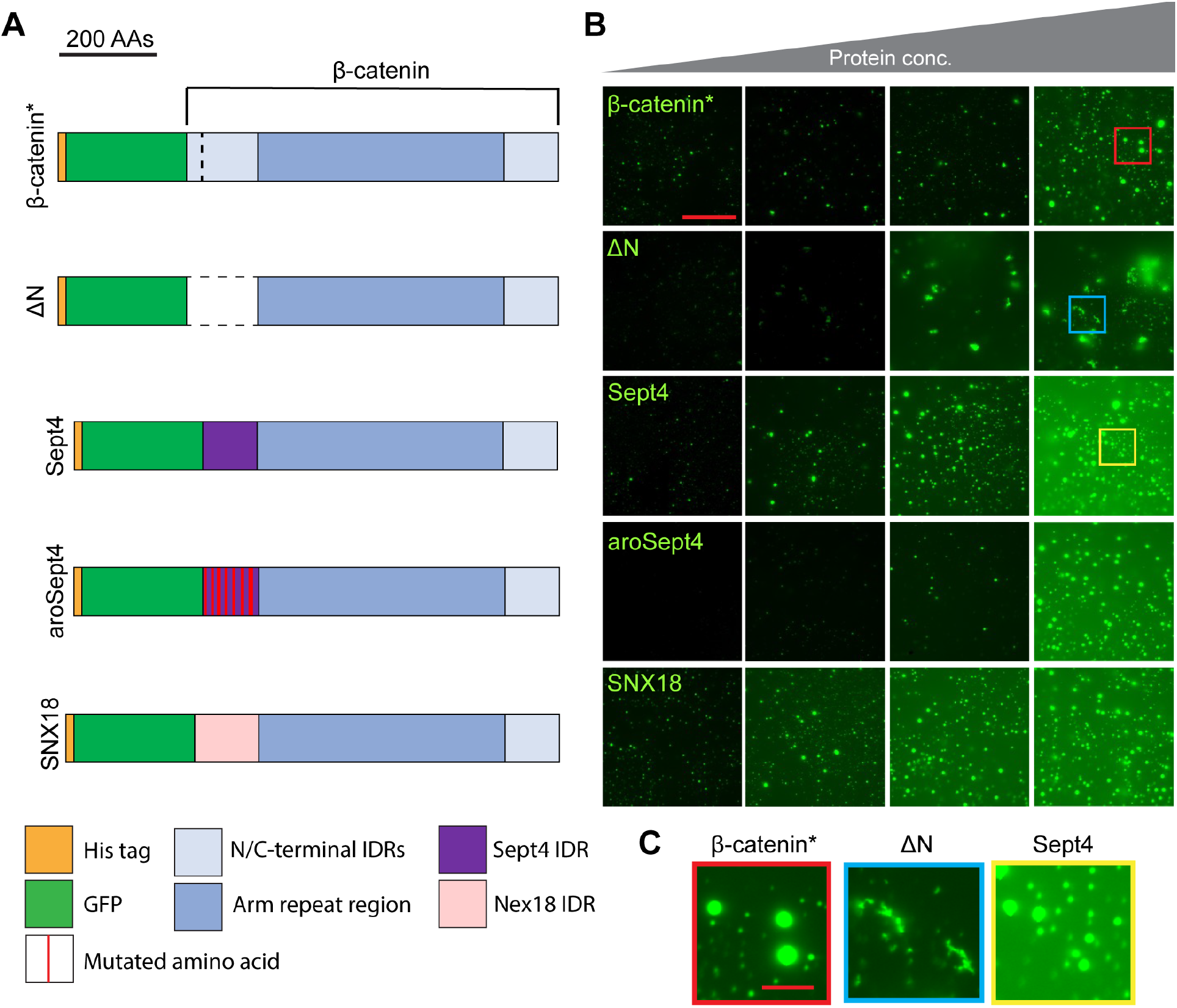
Heterologous IDRs rescue the *in vitro* droplet formation of an N-terminal β-catenin deletion mutant. (A) Cartoon representation of the eGFP-β-catenin* protein, ΔN (amino acids 1-151 deleted), Sept4 (119 amino acids from the Septin4 IDR), aroSept4 (Sept4 IDR with 11 aromatic amino acid mutations), and SNX18 (139 amino acids from the SNX18 IDR). (B) Representative images from an *in vitro* droplet formation assay with the indicated mutants. A protein concentration series of approximately 2μM, 3μM, 4μM, and 8μM is depicted. Droplet assays were performed in 300mM NaCl and 10% PEG-8000. At lower concentrations, ΔN is deficient in droplet formation; at higher concentrations ΔN forms non-spherical structures. The Sept4 and SNX18 droplets are qualitatively similar to those of β-catenin*, while aroSept4 is deficient in droplet formation compared to Sept4. Scale bar = 20μm. (C) Insets from the highlighted regions of panel B.

We wanted to test if these heterologous IDR β-catenin mutants could rescue transcriptional activity of β-catenin. We utilized the TopFlash-Luciferase transcriptional reporter in a HEK293T β-catenin KO cell line [52]. We transiently transfected these cells with the reporter and expression constructs for the heterologous IDR β-catenin mutants (Fig 9A). The ΔN construct is deficient in activity relative to β-catenin*. The Sept4 and SNX18 heterologous IDRs rescue transcriptional activity and mutating the aromatic residues within the Sept4 IDR (aroSept4) ablates activity, again in a manner reminiscent of aroN. The differences in reporter activity are not due to differences in expression (Fig 9B). Additionally, we made the equivalent heterologous IDR mutants in Arm and tested the activity of Sept4-Arm in the fly eye (Fig 9C and S11 Fig for protein sequences). Sept4-Arm has an intermediate effect on eye development, leading to an eye that is approximately 20% smaller than a fly eye which is overexpressing the ΔN-Arm mutant (Fig 9D). Mutating the aromatic residues within the Sept4 IDR returns the level of Arm activity back to ΔN-Arm levels. Our data shows that heterologous IDRs, which rescue β-catenin BMC formation *in vitro,* can also rescue transcriptional activity *in vivo*. Additionally, aromatic residues within the heterologous IDRs that are responsible for facilitating this activity, providing strong evidence for a model where the aromatic residues within β-catenin’s IDRs are important for transcriptional activity through a biomolecular condensation mechanism.

**Fig 9.**
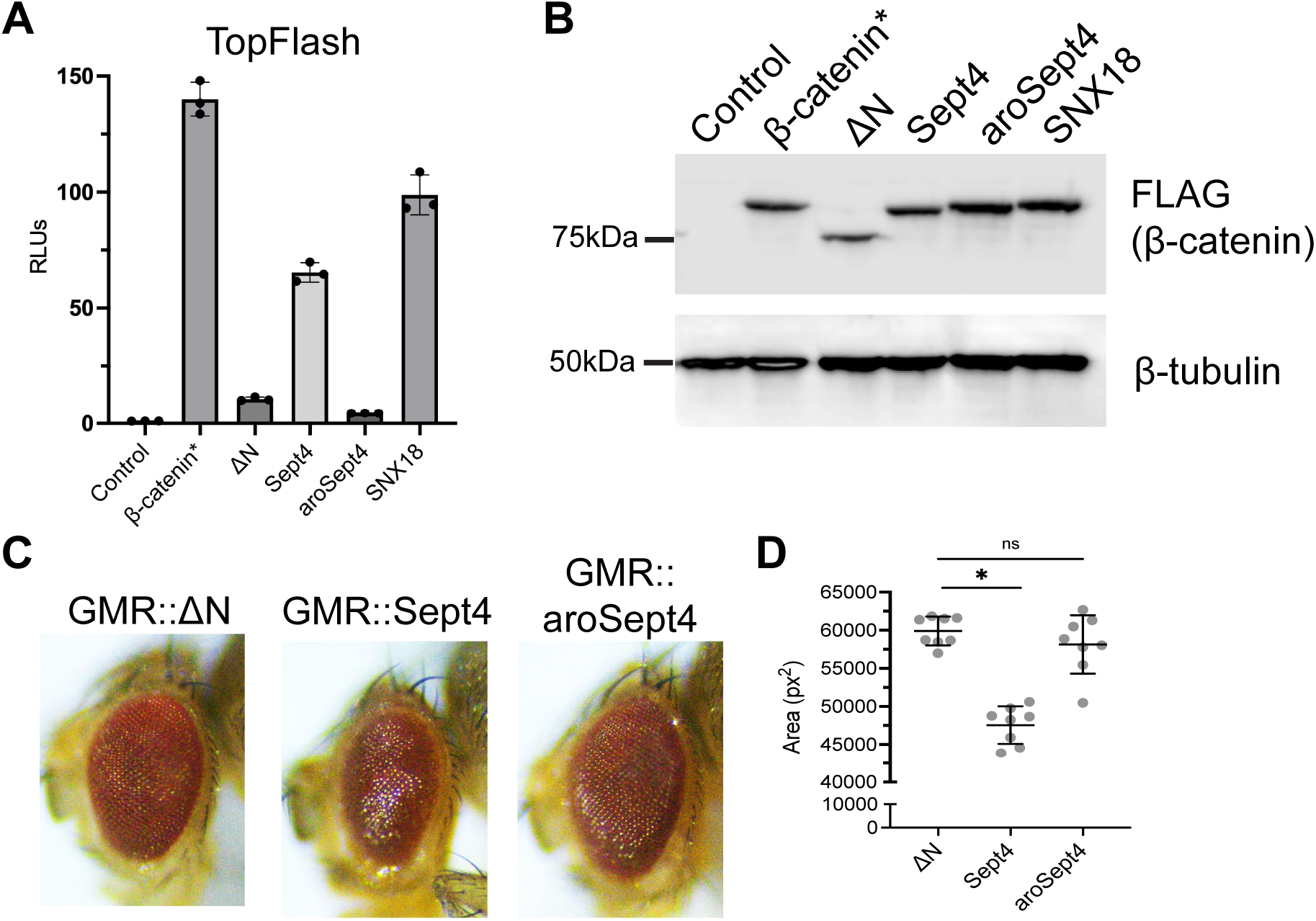
Heterologous IDRs can rescue the activity of a N-terminal β-catenin deletion mutant. (A) TopFlash luciferase reporter activity induced by β-catenin* or the derivative mutant constructs in HEK293T β-catenin KO cells. Cells were transfected with separate plasmids encoding the reporter gene and the FLAG-β-catenin mutant constructs. Luciferase activity was assayed 24hr post-transfection. Data are plotted as mean ± SD (n = 3). (B) Western blot analysis of transfected HEK293T β-catenin KO cell lysate. Lysate samples correspond to the luciferase reporter data. α-FLAG blot shows β-catenin expression. α-β-tubulin was used as a loading control. (C) Micrographs of adult Drosophila eyes containing P[GMR-Gal4] and UAS lines expressing ΔN, Sept or aroSept4. (D) Quantification of adult Drosophila eye area. Data are presented as mean ± SD (n = 8). p-values were calculated by one-way ANOVA followed by Dunnett’s test. * = p < 0.05 and ns = p > 0.05.

## Discussion

This study provides evidence that β-catenin’s ability to form BMCs is essential for its function as a transcriptional co-regulator. Mutations in β-catenin that affect the ability to form BMCs *in vitro* correlate with reduced activity as a transcriptional co-regulator cultured human cells (Fig 1-3). This correlation was also observed *Drosophila*, using several functional readouts, such as transcriptional reporters and developmental phenotypes (Fig 3 and 7). The finding that substitution of specific residues in the N-IDR (or deletion of the N-IDR) severely compromised BMC formation and signaling activity is crucial to our argument, as the N-IDR hasn’t been thought to play a role in transcriptional activation [50,53]. Building on this result, the most compelling evidence linking BMCs to β-catenin’s function involved replacing the N-IDR of β-catenin with two heterologous IDRs from proteins with no known role in transcription. These chimeric β-catenins rescue the deficiencies of the N-IDR deletion in BMC formation and provide significant rescue in transcriptional regulation (Fig 8 and 9). Taken together, our data provide strong support for a model where the ability of β-catenin to form BMCs is an important mechanism for its function as a transcriptional co-regulator.

The concept of BMCs playing an important role in transcriptional activation has generated a large level of support [15,16,19,54] but it is not without controversy. The Mediator subunit MED1 readily forms BMCs with Pol II subunits *in vitro*, and dynamic puncta containing both complexes can be visualized on regulatory chromatin in cultured cells [55]. MED1 and co-activators such as p300 and BRD4 also form mixed BMCs with various TFs *in vitro* and *in vivo* [16,56–59]. Further studies link Pol II transcriptional bursting to a BMC model of regulatory control [19,55,60]. However, live imaging studies linking droplet formation with increased transcriptional output have produced conflicting results [61,62]. Genetic evidence linking the ability of co-activators to undergo phase separation and perform its function in transcriptional activation are limited [56,63]. There is a pressing need for further genetic studies in physiologically relevant contexts to Probe the role of condensates in gene regulation.

Our work builds upon the work of Zamudio and colleagues [20] by providing an extensive functional characterization of β-catenin mutants that are deficient in BMC formation. We find that the observed deficits in β-catenin’s function cannot be attributed to a defect in nuclear import (Fig 3). Considering this, the ChIP-seq data in Zamudio and colleagues support a model in which β-catenin recruitment at WRE chromatin is driven by a combination of the Arm repeats (presumably due to direct binding to TCFs) and IDRs (presumably allowing β-catenin to be enriched in condensates on WRE chromatin). Our heterotypic *in vitro* droplet formation assay data with β-catenin and LEF1 corroborates this idea. β-catenin lacking both IDRs, which is unable to form homotypic condensates (Fig 1) can still be recruited to LEF1 BMCs, although at a reduced level compared to full length β-catenin (Fig 2). We suggest that IDR-driven condensation acts as an amplifier for protein-protein interactions between structured domains, allowing a sufficient concentration of co-activators to facilitate transcription.

### Forces driving β-catenin condensate formation

One commonly proposed mechanism for BMC formation invokes pi-pi interactions between the side chain of aromatic amino acids [64]. As previously reported [20] and extended in this report, aromatic residues in the terminal IDRs of β-catenin play a key role in the ability to form condensates *in vitro* and *in vivo*. Mutation of these residues also had a context-dependent effect on β-catenin’s ability to activate Wnt targets. To narrow down the number of mutations (aroN has 9; aroC 10) we mutated subsets of aromatic residues inspired by the sticker-spacer model [35] and tested for activation of a luciferase reporter. Our data indicated that both “sticker” and “spacer” aromatic residues were crucial for β-catenin activity (S4 Fig). Comparison of pan-aromatic and conserved (between flies and humans) aromatic amino acids in *Drosophila* developmental assays (Fig 5 and 6) suggest that the conserved residues (7 in the N-IDR; 5 in the C-IDR) might be the most important. In *Drosophila* transgenic assays, we compared the effect of mutating all aromatics (as with human β-catenin, 9 in the N-IDR and 10 in the C-IDR) with only the residues conserved between flies and humans (7 in the N-IDR; 5 in the C-IDR). As the pan-aromatic and conserved mutants had similar defects in signaling (Fig 5 and 6), it is tempting to suggest that the conserved residues contribute most to Arm/β-catenin’s activity. The N- and C-IDRs contain a mixture of tyrosines, phenylalanines and tryptophans (S2 and S7 Fig). BMC formation of some proteins, for example, FUS, are predominately driven by tyrosine and arginine interactions. Mutating tyrosine residues to phenylalanine strongly reduces the ability of FUS to form BMCs [65]. However, some of our mutants containing multiple phenylalanine substitutions, e.g., N-sticker (3 of 5) and aroN-con (5 of 7) have severe signaling defects (Fig 5 and 6, S4 Fig), suggesting that these aromatic residues are important for condensation/activity. Our data is consistent with the idea that BMC formation is driven by multivalent interactions of all three types of aromatic residues and that many/most of the 19 residues in the IDRs contribute condensation and transcriptional activity. However, further mutagenesis is needed to test the relative contributions of each aromatic position.

### Is β-catenin condensation universally required for Wnt target gene activation?

Our data support an important role for β-catenin condensation in transcriptional activation, but it is unclear if this is a universal requirement for the expression of all Wnt target genes. In nearly every case we examined (the one exception being aroN activating Axin2 in Hela cells; Fig. 4A), mutation of aromatic residues resulted in a reduction of signaling activity. However, the degree of this defect depended on the assay employed. For the reporter gene assays in human cells (Fig 2, S4 Fig) and the developing *Drosophil*a eye there was a strict requirement for aromatic residues, e.g., the five substitutions in aroC-con abolished Arm’s signaling activity in the eye (Fig 5). For endogenous targets in Hela cells (Fig 4) and a Wg/Wnt reporter in wing imaginal discs (Fig 6), there was an intermediate defect in signaling. In the *Drosophila* embryo, a mutant (aroNC) with 19 substitutions still had the ability to rescue a strong *arm* loss-of-function phenotype (Fig 7). It is possible that some of this rescue is the result of aroNC de-repressing Wnt target gene expression through displacing co-repressors from TCF/Pangolin [66,67], but the modest level of expression of the aroNC transgene makes this unlikely. Clouding the interpretation is the fact that β-catenin lacking all 19 aromatic amino acids retains the ability to form BMCs *in vitro* at high concentrations (Fig 1). This is different from the results reported by Zamudio et al., but we note that in this study, the GFP-b-catenin fusions were purified under denaturing conditions and then renatured (REF). This raises the possibility that aroNC did not properly refold. Given our results that aroNC can form BMCs (albeit only at higher concentrations), further studies are needed to identify additional residues in the N- and C-IDRs that contribute to condensate formation. This would allow the construction of a tighter condensate mutant, to address whether BMC formation is universally required for activation of Wnt targets with increased certainty.

### Non-transcriptional BMCs containing β-catenin

In the absence of Wnt stimulation, β-catenin is targeted for proteasomal degradation by a “destruction” complex consisting of AXIN, APC, the kinases CKI and GSK3, and the E3-ubiquitin ligase Tr-BP [68]. The multivalency of the protein-protein interactions between destruction complex members led to the suggestion that it formed a BMC [68]. Indeed, the ability of AXIN to undergo phase separation has been genetically linked to efficient down-regulation of β-catenin [69] and evidence for a destruction complex BMC at endogenous levels of expression has been reported [70]. The positive Wnt effector Dvl2 has also been shown to form condensates, which has been suggested to play a role in inhibiting the destruction complex [71–73]. The aromatic β-catenin/Arm mutants described in this report also contained point mutations blocking GSK3 phosphorylation, rendering them insensitive to degradation by the destruction complex. This allowed their role in transcriptional regulation to be unambiguously assayed. Further studies are needed to determine whether β-catenin mutants such as aroNC can be efficiently recruited to the destruction complex.

### Conclusions

Our data suggests that there is a high degree of context-dependency regarding the relationship between aromatic/condensation and activation of specific Wnt targets. Understanding the molecular basis for this specificity will require a combination of transcriptomics and an in-depth examination of WREs that have a strong requirement for aromatic residues in N-IDR and C-IDR and those that do not. Nonetheless, this report provides strong evidence that a role for β-catenin condensation needs to be considered to fully understand how the Wnt pathway activates transcription.

## Materials and Methods

### Plasmids

FLAG-tagged β-catenin variants were expressed in transient transfection assays using the pCDNA3.1 vector (Thermo Fisher Scientific). A plasmid expressing human β-catenin containing a S33Y mutation (pCDNA3-S33Y) was the starting point for further mutagenesis [74]. β-catenin mutants were created using gBLOCKS (Integrated DNA Technologies) which were subcloned into pCDNA3-S33Y that was linearized with either *BamHI* & *PmlI* (N-IDR mutants) or *BbvCI* & *XbaI* (C-IDR mutants). To express the β-catenin proteins in *E. coli*, a pET28 expression vector expressing a His-tagged GFP-β-catenin (RY8686), which was a gift from R. Young (MIT) was used [20]. Various β-catenin variants were subcloned into RY8686 using *BamHI* & *NotI*.

For the luciferase assays, the Topflash and CREAX reporter plasmids were constructed using pGL4.23 (Promega). Specific transcription factor binding sites or regulatory elements are upstream of a minimal TATA-box promoter driving the expression of the firefly luciferase gene. TopFlash has 6x TCF binding sites (plasmid was a gift from E. Fearon, University of Michigan). The CREAX luciferase reporters contain the endogenous WREs from the human *Axin2* locus [33]. For the *Defa5* reporter, the Defa5 promoter (WRE plus promixal promoter) was cloned into the promoterless pGL4.10 plasmid [34].

The pCW57.1 vector (gift from David Root, addgene plasmid #41393) was used to generate the lentiviral particles for cell transduction. In brief, the coding regions of various Flag-tagged β-catenin constructs were PCR amplified from the aforementioned pCDNA3 vectors, along with the SV40 polyadenylation site. Overlapping sequence was included to allow these amplicons to be combined with a 7.8kB *NdeI*/*SalI* fragment of the pCW57.1 lentiviral vector via Gibson assembly using the NEBuilder HiFi DNA Assembly Master Mix (New England Biolabs). Products were confirmed with Sanger sequencing (Genewiz, South Plainfield, NJ).

Vectors for transgenic *Drosophila* strains were constructed using the PhiC31 transgenesis system [40]. An *arm* cDNA encoding two activating mutations (T52A and S56A) was cloned into a pUAST-FLAG-attB vector (a gift from CY Lee, University of Michigan). This construct (pUAST-Arm*-FLAG-attB) expresses the stabilized Arm protein tagged with a C-terminal FLAG epitope. Additional Arm mutants were created using gBLOCKS (Integrate DNA Technologies) cloned into pUAST-Arm*-FLAG-attB using either *MluI* & *StuI* (for N-terminal IDR mutants) or SacI & ClaI for C-terminal IDR mutants.

### Protein Purification

pET28 vectors encoding the various β-catenin mutants were transformed into C41 cells and plated on LB plates containing kanamycin (50μg/ml). Multiple fresh bacterial colonies were inoculated into 5ml LB broth with kanamycin and grown overnight at 37°C. The overnight cultures were diluted in 500ml of fresh LB broth with kanamycin and grown at 37°C until the cultures reached an OD_600_ of 0.6-0.8. IPTG was added to a working concentration of 1mM, and the cells were grown for 18 hours at 16°C. Cells were collected by centrifugation.

Pellets from the 500ml cultures were resuspended in lysis buffer (50mM Tris-HCl pH: 7.5, 300mM NaCl, 4mM imidazole, 0.1% Triton-X 100, 1 tablet complete protease inhibitors (Roche, 11873580001)). Samples were then sonicated, 10 seconds on, 20 seconds off, for 3 minutes. Lysates were then cleared by centrifugation at 12,000xg for 12 minutes and added to 1ml of pre-equilibrated TALON metal affinity resin (Takara Bio, 635502). The slurry was rotated at 4°C for 1 hour. The slurry was then centrifuged at 700g for 5 minutes at 4°C. The bead pellets were then washed twice with 10ml of wash buffer (50mM Tris-HCl pH: 7.5, 300mM NaCl, 8mM imidazole). Protein was eluted in 1.5ml of elution buffer (50mM Tris-HCl pH: 7.5, 300mM NaCl, 50mM imidazole, 10% glycerol) and the samples were rotated for 10mins at 4°C. Protein samples were then concentrated using Amicon Ultra centrifugal filters (30k MWCO, Millipore). The protein concentration of eluates were estimated with a Bradford assay (BioRad) and analyzed on an 8.5% acrylamide gel stained with Coomassie.

### *In vitro* droplet formation assay

Assays were carried out on chambered coverglass slides (Grace Bio-Labs, 112359) that were passivated with Pluronic F-127 (Sigma-Aldrich, P2243). A 5% (w/v) Pluronic F-127 solution was added to the slide’s chambers and incubated for at least 1 hour. After the incubation, the chambers were washed with buffer (50mM Tris-HCl pH: 7.5, 300mM NaCl, 10% glycerol). Varying concentrations of eGFP- and mCherry-tagged proteins were added to the chambers with droplet formation buffer (50mM Tris-HCl pH: 7.5, 300mM NaCl, 10% glycerol, 10% PEG-8000). The mixture was incubated for 15 minutes before being imaged on a Leica DMI6000B with a 63x objective and a Hamamatsu ORCA-R2 camera. Images were processed and analyzed using Leica Application Suite X (LAS X). Line profile intensities were calculated using a single line through representative condensates.

For the hexanediol sensitivity assays, 10% (w/v) 1,6-hexanediol and 2,5-hexanediol (Sigma-Aldrich, 240117 and H11904, respectively) solutions were made. These solutions were added to protein diluted in buffer (50mM Tris-HCl pH: 7.5, 300mM NaCl, 10% glycerol) before or after PEG-8000. All droplet assays shown were repeated at least three times on separate days (often with separate protein preps) and the results shown are representative.

### Cell Culture, transfection, stable cell line generation, DOX treatment

HEK293T cells were obtained from the American Type Culture Collection. HeLa cells were a gift from Y. Wang (University of Michigan) and HEK293T β-catenin KO cells were a gift from K. Basler (University of Zurich) [52]. All cells were grown at 37°C with 5% CO_2_ and in Dulbecco’s modified Eagle’s medium (Gibco, 11995065) supplemented with 10% fetal bovine serum and penicillin-streptomycin-glutamine (Gibco, 10378016).

For transfection in HEK293T cells, 50,000 cells per well were plated in a 48 well plate and grown overnight. Cells were transfected using polyethylenimine-MAX (PEI-MAX, PolySciences, 24765-1) following the manufacturer’s protocol. All luciferase assays were performed at least three times on separate days, with similar results obtained in each experiment.

For this study, stable HeLa cell lines containing DOX-inducible β-catenin mutant (β-catenin*, aroN, aroC, and aroNC) expression cassettes were generated by lentiviral transduction. Lentiviral supernatants were made by the University of Michigan Vector core lab. To generate the mutants, HeLa cells were incubated with viral supernatants for 24 hours, then the cell culture medium was replaced with fresh medium and cells were grown for an additional 24 hours. Transduced cells were then selected for and maintained in cell culture medium containing 1μg/ml puromycin. For DOX-induced expression of the mutant proteins, the individual HeLa cell lines were treated with varying doses of DOX to normalize expression across the tested mutants.

### Western Blotting

Cell samples were lysed and denatured in hot 1x SDS loading buffer. Protein samples were separated by SDS-PAGE and transferred to a polyvinylidene fluoride membrane (PVDF, Bio-Rad, 1620177) and blocked in 5% bovine serum albumin (BSA). Protein blots were incubated in primary antibody (diluted in 5% BSA) overnight at 4°C. After the incubation, protein blots were washed three times with tris-buffered saline containing 1% Tween-20 (TBS-T), then incubated with a secondary antibody (diluted in 5% BSA) for 1 hour at room temperature. Blots were then washed three times with TBS-T, developed with a chemiluminescent substrate (Pierce, 34577), and imaged using a LI-COR Odyssey CLx. Images were processed in Adobe Photoshop.

Antibodies used: anti-FLAG-horseradish peroxidase (HRP, Sigma-Aldrich, A8592, 1:5000), anti-β-tubulin (Proteintech, 66240-1, 1:20,000), anti-mouse HRP (Jackson ImmunoResearch, 115-035-003, 1:2000).

### Immunofluorescence

HEK293T and HeLa cells were seeded on 12mm round glass coverslips (Warner Instruments, 64-0712) in 24 well plates and grown overnight. The following day, cells were either transfected or treated with DOX to express FLAG-tagged β-catenin mutant proteins. 24 hours after transfection or DOX treatment, cells were fixed with 4% paraformaldehyde (Electron Microscopy Sciences, 15710). The IF protocol was previously published (Leica, Quick guide to STED sample preparation). Briefly: following fixation, the cells were washed with PBS and permeabilized with 0.1% Triton-X 100 (MP Biomedicals, 807426). Cells were blocked with 4% bovine serum albumin for 1hr, then incubated in primary antibody solution overnight at 4°C. The following morning, cells were washed and incubated in secondary antibody solution for 1 hour at room temperature. Cells were then washed and counterstained with DAPI. Coverslips were mounted on slides with Vectashield Mounting Media (Vector Labs, H-1000). Images were acquired with a Leica Sp8 laser confocal microscope and processed using LAS X.

Antibodies used: anti-FLAG (Sigma-Aldrich, F3165, 1:1000), anti-mouse-Alexa568 (Molecular Probes, A11031, 1:1000), anti-Cut (Developmental Studies Hybridoma Bank, 2B10, 1:20)

### qRT-PCR

Total RNA was extracted using the Rneasy Plus Mini Kit (QIAGEN, 74134). cDNA synthesis was done using SuperScript III reverse transcriptase (Invitrogen, 18080-044) with oligo-dT primers. For the qRT-PCR, *Power*SYBR Green PCR Master Mix (Applied Biosystems, 4367659) was used and the reaction was carried out in a StepOnePlus Real-Time PCR System (Applied Biosystems). The β-actin and 18s genes were used as internal controls, and the relative expression of target genes was calculated using a modified Pfaffl equation which accounts for multiple reference genes [75,76]. Primer sequences listed in table S1. Experiments were repeated three times with qualitatively similar results obtained.

### Transgenic *Drosophila* strains

pUAST-Arm*-FLAG-attB and other UAS-arm derivatives were injected into M{3xP3-RFP.attP’}ZH-51C and M{3xP3-RFP.attP}ZH-86Fb embryos by BestGene Inc (Chino Hills, CA) or Rainbow Transgenic Flies (Camarillo, CA). Transformants were identified by the presence of the mini white gene. Transgenic chromosomes were balanced over the SM5a-TM6B compound balancer, either as single inserts or in combination (51C & 86Fb). Insertions at 51C were meiotically recombined with the P[GMR-Gal4] transgene. Other Gal4 lines were obtained from the Bloomington Stock Center. Da-Gal4 and Arm-Gal4 were meiotically recombined onto a single third chromosome and balanced over TM6c. All *Drosophila* stocks were raised on yeast/glucose food and experiments were performed at 25°C unless otherwise indicated.

### Imaging of Drosophila eye and wing tissues

Adult flies containing P[GMR-Gal4] and four copies of P[UAS-arm*] or its variants were frozen at −20°C overnight and photographed with a Leica Stereo Dissecting Scope (Leica DMI6000B) attached to a digital camera. Eye size was quantified using ImageJ. Crosses were repeated multiple times with similar results. For Cut immunostaining, white prepupa were selected and aged 40-44 hrs at 25°C before dissection and fixation with 4% paraformaldehyde. Pupal eyes were stained with mouse anti-Cut (Developmental Studies Hybridoma Bank; 1:100). At least ten eyes were examined for each condition, with similar results.

Wg/Wnt signaling in the wing imaginal discs was measuring using a Wnt GFP synthetic reporter previously described [45]. This reporter contains three copies of a grainy head binding site and four copies of a TCF-Helper upstream of a minimal promoter driving GFP. Larva containing this reporter and one copy each of P[Dpp-Gal4] and P[Tub-Gal80ts] and a single copy of P[UAS-arm*] or its variants were reared at 18°C and then shifted to 29°C for 18 hours before selecting late third larval instar for dissection, fixation with 4% paraformaldehyde and mounting. Crosses were repeated multiple times and at least twelve discs were visualized for each condition.

Quantification of fluorescent reporter activity was performed in ImageJ. A region of interest (ROI) was defined in the location of ectopic reporter activity, integrated density of the fluorescent signal was quantified, and the same ROI and calculation was used at a site of endogenous reporter activity. Data is presented as a ratio of the integrated density of the reporter at the ectopic activation site to the endogenous activation site.

To monitor expression of Arm* proteins, wing discs treated as described above and eye/antennal discs were fixed and subjected to IF using a mouse anti-FLAG antibody (Sigma-Aldrich, F3165, 1:100) and anti-mouse-Alexa568 (Molecular Probes, A11031, 1:200) as previously described [77]. All GFP and IF images were acquired with a Leica SP8 laser confocal microscope and processed using LAS X.

### Preparation of embryonic *Drosophila* cuticles

Embryonic *Drosophila* cuticles were prepped for imaging using a previously described method [78]. Briefly, grape agar plates were added to *Drosophila* cultures for egg collection. Eggs were incubated at 25°C and allowed to develop to the point of death, which occurs in late embryogenesis.

The embryos were dechorionated by placing them in a 50% bleach solution for 2 minutes, then rinsing them with distilled water, and then dried. The embryos were then devitellinized by transferring them to a 1:1 heptane to methanol solution, and vigorously vortexing for 30 seconds. The heptane and methanol solution was decanted and the embryos were washed 3 times with methanol and a final time with a 0.1% Triton-X 100 in methanol solution. The embryos were then transferred to a glass slide, residual methanol was evaporated, and the embryos were mounted in a 1:1 solution of Hoyer’s mounting medium (Hempstead Halide) and lactic acid. The embryos were imaged using a Nikon E800 26prightt microscope equipped with a Nikon DS-Fi3 camera and a Nikon Dark Field Condenser (Dry 0.95-0.80). Images were processed using NIS-Elements software.

Various P[UAS-arm*] strains were crossed with P[Da-Gal4],P[Arm-Gal4] which are both active throughout the embryonic epidermis [45,46]. Embryos contained one copy of each P[Gal4] and two copies of P[UAS-arm] transgenes for the experiments described in Fig. 6. To test for rescuing activity of the different *arm* transgenes, males homozygous for P[UAS-arm] transgenes were crossed to *arm^4^*/FM7; P[Da-Gal4] females. *Arm^4^*is an amorphic allele of arm that produces a protein truncated in the sixth Arm repeat that produces no detectable protein [79]. Approximately 3/4s of the progeny displayed a consistent phenotype indistinguishable from those of embryos with P[Da-Gal4] and the respective P[UAS-arm] construct. Approximately one quarter had a highly penetrant but distinct phenotype consistent with an *arm^4^*/Y embryos with significant phenotypic rescue. All crosses were repeated multiple times; the phenotypes obtained were highly penetrant (n>20 for each condition).

## Acknowledgements

The authors would like to thank Richard Young and his co-workers for inspiring this study, and for providing their GFP-β-catenin plasmid. Thanks to Claudio Cantù and Konrad Basler for providing the HEK293T β-catenin knockout cell line. Special thanks to Sarah Bui for assistance with lentivirus infections and line analysis of heterotypic BMCs, and to Yanzhuang Wang for use of his BL2 biological safety cabinet. Thanks to Anthony Vecchiarelli for discussions on biomolecular condensates. Thanks to Hwajeong Yi, Jon Millar, Jonathan Calderon Juarez and Carla Peralta for assistance with the construction of plasmids and to Aravind Ramakrishnan and Jon Millar for critical reading of the manuscript.

